# Forward simulations of walking on a variable surface-impedance treadmill: A comparison of two methods

**DOI:** 10.1101/2021.10.11.463993

**Authors:** Banu Abdikadirova, Mark Price, Wouter Hoogkamer, Meghan E. Huber

## Abstract

Recent experiments with a variable stiffness treadmill (VST) suggest that modulating foot-ground contact dynamics during walking may offer an effective new paradigm for gait rehabilitation. How gait adapts to extended perturbations of asymmetrical surface stiffness is still an open question. In this study, we simulated human gait with prolonged asymmetrical changes in ground stiffness using two methods: (1) forward simulation of a muscle-reflex model and (2) optimal control via direct collocation. Simulation results showed that both models could competently describe the biomechanical trends observed in human experiments with a VST which altered the walking surface stiffness for one step. In addition, the simulations revealed important considerations for future experiments studying the effect of asymmetric ground stiffness on gait behavior. With the muscle-reflex model, we observed that although subtle, there was a difference between gait biomechanics before and after the prolonged asymmetric stiffness perturbation, showing the behavioral signature of an aftereffect despite the lack of supraspinal control in the model. In addition, the optimal control simulations showed that damping has a large effect on the overall lower-body muscle activity, with the muscle effort cost function used to optimize the biomechanics increasing 203% between 5 Ns/m and 2000 Ns/m at a stiffness of 10 kN/m. Overall, these findings point to new insights and considerations for advancing our understanding of human neuromotor control of locomotion and enhancing robot-aided gait rehabilitation.

## I. INTRODUCTION

Locomotion is one of the most important aspects of human mobility and any type of locomotion dysfunction can greatly affect someone’s quality of life. Traditional gait therapies typically require the constant presence of a therapist which makes them expensive and inefficient. To mitigate these problems, significant advancements in the development of robotic rehabilitation devices have been made in the last decade which could help gait rehabilitation become more autonomous and independent of a therapist. At the moment, however, robot-aided gait rehabilitation is still not as effective as the current standard of care [1], [2].

Most gait rehabilitation robots operate by applying torques directly to the wearer’s joints. The effect of controlling the mechanical impedance (the dynamic generalization of stiffness) of the foot-ground interface to alter the gait behavior has been relatively unexplored. Foot-ground mechanical impedance strongly influences the shape and magnitude of the ground reaction force profile, which influences the overall biomechanics. For example, the magnitude of ground reaction forces has been shown to change ankle range of motion and ankle dorsiflexor m uscle f orces [3]. Additionally, foot-ground interaction dynamics directly affect foot stability, which is a crucial indicator of body balance overall [3]. The fact that gait behavior can be modulated by altering foot-ground interaction dynamics suggests that it is a promising approach for gait therapy. However, there is relatively little knowledge as to how human gait responds to perturbations of foot-ground stiffness over time and whether such behavioral changes arise from neuromotor adaptation and/or caused by biomechanical effects [4].

To address this gap, two research groups have developed variable stiffness treadmills (VST) which can change the stiffness of the walking surface in real time [5], [6], allowing experimental study of gait changes in response to stiffness perturbations and the effectiveness of the approach as a rehabilitation tool. Skidmore and Artemiadis [7] investigated the effect of walking surface stiffness on inter-leg coordination by applying single-step asymmetric low stiffness perturbations of various magnitudes. The results of their study indicate increased activation of TA and SOL muscles, as well as increased flexion of all three lower-limb joints in the unperturbed leg of subjects with decreasing stiffness perturbation magnitude. A similar set of experiments was performed by the same research group to evaluate whether the brain is involved in inter-leg coordination [8]. EEG activation measurements indicated significant changes in the medial side of the left brain with applied asymmetric low stiffness perturbations which depicts the involvement of supraspinal neural circuitry. The study suggested a strong potential of VST for gait impairment created by stroke where the brain is the root cause of the gait dysfunction. Overall, this kind of novel device provides new insight and holds promising potential in gait rehabilitation.

In support of experiments with a VST, a model-based simulation approach can provide complementary insight as well as create testable predictions for subsequent experiments. To this end, Chambers and Artemiadis [9] performed model-based analysis of asymmetric stiffness perturbations using the same conditions as in human experiments [7]. They implemented a three-dimensional neuromuscular model that has both supraspinal and spinal control layers [10], [11]. The supraspinal control layer regulates two variables considered essential to achieve stable walking: sagittal angle of attack and hip to ankle span. High level controllers were added to the supraspinal layer to correct for model failure after more extreme perturbations. A key limitation of these experiments and simulations, however, is that they only investigate asymmetric ground stiffness perturbations for a single step. To our knowledge, neither the measured nor simulated effect of long-term walking on asymmetric ground stiffness has been reported. Addressing this knowledge gap is critical as adaptation to a persistent perturbation is a key element in other gait rehabilitation techniques such as split-belt treadmill interventions [12]. Given the difficulty of achieving stable walking for a muscle-reflex model with spinal control for a single step perturbation, it is likely that achieving stable walking during persistent ground stiffness asymmetry will similarly require substantial modifications to the approach.

An optimal control simulation methodology may be an alternative for simulating stable, periodic gait with severe ground stiffness asymmetry. Optimal control simulations of human movement have been used to generate gait patterns with close agreement to experimental biomechanics, typically by solving for muscle excitations which minimize some measure for muscle effort [13], [14]. This approach is based on the observation that humans tend to select walking speeds and stride frequencies that minimize the metabolic cost of transport [15], [16]. Importantly for the asymmetric ground stiffness perturbation problem, the simulated gait biomechanics are the solution to an optimization problem which is typically constrained to prevent falling (i.e., the pelvis height must remain above a minimum value) and to enforce periodicity (i.e., muscle states and joint kinematics are identical at the start and end of the gait cycle).

In this study, we implemented an existing muscle-reflex gait model and developed an optimal control gait model to simulate walking on a VST across a range of asymmetricc stiffness perturbations. Our objective was to (1) compare how the two gait models respond to asymmetric changes in ground stiffness and explain the ways in which they agree and disagree, (2) determine whether either model could competently describe the results observed in human experiments [7], and (3) gain insight in the form of testable predictions as to how gait can be altered by modulating foot-ground mechanical impedance for rehabilitation purposes.

## II. METHODS

### A. Muscle-Reflex Model of Locomotion

#### 1) Model Description

The muscle-reflex model of Geyer and Herr [17] is a 2D locomotion model that more accurately describes human walking compared to previously developed simplified bipedal spring-mass model [18]. The model consists of seven segments: one trunk segment, and thigh, shank, and foot segments for each leg (Fig. 1a). Each of the legs has seven Hill-type muscles. Periodic, human-like walking is generated by muscle control signals resembling reflex responses generated by the peripheral nervous system. Each foot has two contact points with the ground. The foot-ground interaction is modelled as a nonlinear spring-damper system normal to the ground, and static and kinetic friction terms parallel to the ground. The dimensions and mechanical properties of the segments, initial positions of the joints, joint motion limits, muscle control parameters and foot-ground interaction parameters are tunable hyperparameters which determine the ability of the model to achieve steady state walking as well as properties of the resulting gait pattern such as the walking speed.

**Fig. 1.**
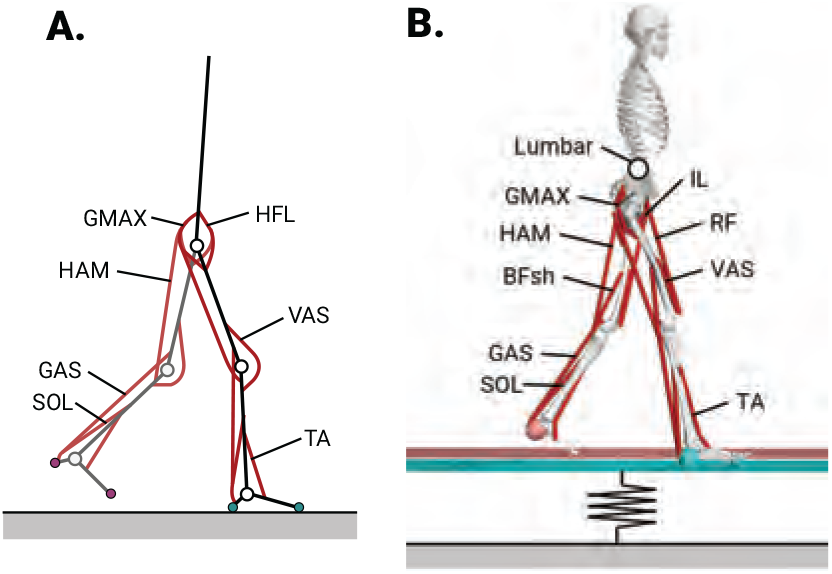
Simulation setup. A) The muscle-reflex model developed by Geyer and Herr [17]. B) The optimal control model with added platforms to simulate a variable stiffness treadmill. Note that this model has two additional muscles per side than the muscle-reflex model (RF, BFsh).

#### 2) Simulation Methods

The muscle-reflex model was simulated in Simulink (MATLAB R2019a, Mathworks, Natick, MA, US). For the sake of simplicity, all the hyper-parameters were kept at their default value except for the ground stiffness under each foot, which was altered to simulate asymmetric ground surface stiffness. The default hyperparameter values are optimized to specifically maintain the average preferred human walking speed (1.2 – 1.4 m/s).

For each simulated trial, the simulation stop time was 120 s which resulted in approximately 90 strides. During the initial 30 strides, the ground under each foot was rigid. More specifically, the rigid stiffness value was 130 kN/m, as it was the highest stiffness value the model could achieve steady-state walking without falling. Over the next 30 strides, the ground stiffness under the right foot was reduced during right leg stance (i.e., when the right foot was in contact with the ground). To match the experimental conditions of [7], ground stiffness under the right foot was set to either 5, 10, 50 or 100 kN/m, while ground under the left foot remained rigid (130kN/m). During the last 30 strides, the ground under both feet was again set to rigid.

### B. Optimal Control Model of Locomotion

#### 1) Model Description

A modified version of a 2D locomotion model implemented in OpenSim “2D gait.osim” [19] to calculate minimum muscle effort optimal control solutions for asymmetric ground surface stiffness. The model is 62.2 kg and 1.64 m tall, and contains 8 body segments and 10 anatomical degrees-of-freedom (DoF). The lower limb joints are actuated by 18 “DeGrooteFregly2016” Hill-type muscles [20] and the lumbar joint is actuated by an ideal coordinate actuator (Fig. 1). This model was chosen as it closely resembles the muscle-reflex model in its simplifications to the muscles, body segments, and contact geometry.

The model was modified by adding two platforms of negligible mass (0.1 kg) connected to the ground with linear spring-damper coordinate actuators, each with a single DoF in the vertical translation axis, to simulate the ideal behavior of the walking surface of a variable stiffness treadmill. Foot-ground contact was modeled using a smoothed Hunt-Crossley viscoelastic contact model [21] combined with a Stribeck friction model [22]. Contact spheres were located at the heel and toe of each foot with flat contact planes located on the top surface of each of the walking platforms, configured to respond only to the corresponding foot. We left this contact model with the parameters from the original model, which approximate the behavior of an athletic shoe [23], and varied the linear spring and damping coefficients of the walking platform coordinate actuators. Access to this modified model will be made available and detailed here at the time of publication.

#### 2) Simulation Methods

The OpenSim model was simulated by solving optimal control problems with a direct collocation approach using Moco 1.0 [24]. A minimum muscle effort cost function was used, and the optimization was constrained to generate periodic strides at 1.2 m/s walking speed (within the range of the baseline muscle-reflex model configuration). The equations of motion and muscle dynamics were enforced as constraints at each of 101 nodes across the span of the gait cycle with a tolerance of 10^*−*4^.

The minimum muscle effort cost function was defined as

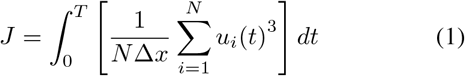

where *u*_*i*_ is the excitation of actuator *i* at time *t, N* is the total number of actuators in the model, and Δ*x* is the horizontal distance traveled by the model over one stride. This cost function approximates muscle fatigue and has been demonstrated to result in realistic kinematics and ground reaction forces for level walking [13].

Because the optimal control simulations were constrained to simulate periodic strides, transient behavior cannot be simulated; each ground stiffness condition is simulated as a stable walking pattern at steady state. Similar to the muscle-reflex model simulations, the ground stiffness under the right foot was set to either 5, 10, 50 or 100 kN/m, while ground under the left foot remained rigid (1000 kN/m). The ground damping coefficient under each foot was set to 1000 Ns/m, which overdamps the response similarly to the default nonlinear damping value in the muscle-reflex model.

In addition, an array of 16 spring constants (5–1000 kN/m) and 7 damping coefficients (5–2000 Ns/m) for a total of 112 unique combinations was simulated. This extended search with the optimal control model was conducted to explore the effect of changing both stiffness and damping on gait well outside of the range of stable walking conditions than can be achieved with the muscle-reflex model.

### C. Comparative Analyses

For each of the four asymmetric ground stiffness conditions, comparisons of the kinematics and muscle activity were made between (1) the two simulated model and (2) the simulated models and the experimental results of [7]. Muscle activity was compared by calculating the mean muscle activation across the gait cycle for each stiffness condition, and normalizing these values against the corresponding mean activations for the symmetrical rigid ground condition. For a subset of muscles, muscle activity curves with respect to the gait cycle were also compared. These curves were normalized against the peak activation of the corresponding muscles for the rigid condition. Note that normalization of the muscle activity results was performed to account for differences in the muscle parameters and model mass parameters.

### D. Additional Model Analyses

The following two analyses could be not be comparative as (1) each additional simulation could only be achieved with one model or the other, not both, and (2) there are no experimental observations available for comparison. Nevertheless, these analyses provide valuable considerations for designing and interpreting the results of future experiments with asymmetric impedance perturbations. Simulation of the muscle-reflex model over time allowed us to assess how gait kinematics change with the application and removal of the asymmetric stiffness perturbation. In addition, simulation of the optimal control model with varying stiffness and damping parameters allowed us to explore how varying the different ground mechanical impedance parameters could be tuned to elicit greater changes in gait behavior.

## III. SIMULATION RESULTS

### A. Muscle-Reflex Model vs. Optimal Control Model Results

The muscle-reflex model was successfully able to reach steady-state walking for ground perturbations with stiffness values of 50 kN/m and 100 kN/m. However, the model failed to achieve stable walking and fell over for stiffness values of 5 kN/m and 10 kN/m after a single stride and 7 strides, respectively. All optimal control simulations converged within the constraint tolerances.

#### 1) Comparison of Kinematics

Fig. 2 represents the steady-state hip, knee, and ankle joint kinematics for both optimal control and muscle-reflex simulations. The overall kinematic profiles are in agreement with each other, however, there are noticeable discrepancies between the two models. The optimal control simulation predicts between 10–30^*°*^more hip flexion on both sides than the muscle-reflex simulation across the gait cycle, indicating some additional forward trunk lean. The muscle-reflex model tends to overestimate the knee flexion angle during stance for both sides, relative to conventional walking and experimental data [7]. The muscle-reflex model also predicted a plantarflexion spike during early stance for both sides, indicative of toe-slap, whereas the optimal control model has a more conventional ankle angle profile. For the optimal control model, kinematic changes from baseline mostly occur for the 10 kN/m stiffness condition or lower, with very little observable difference between the 100 kN/m and 50 kN/m conditions.

**Fig. 2.**
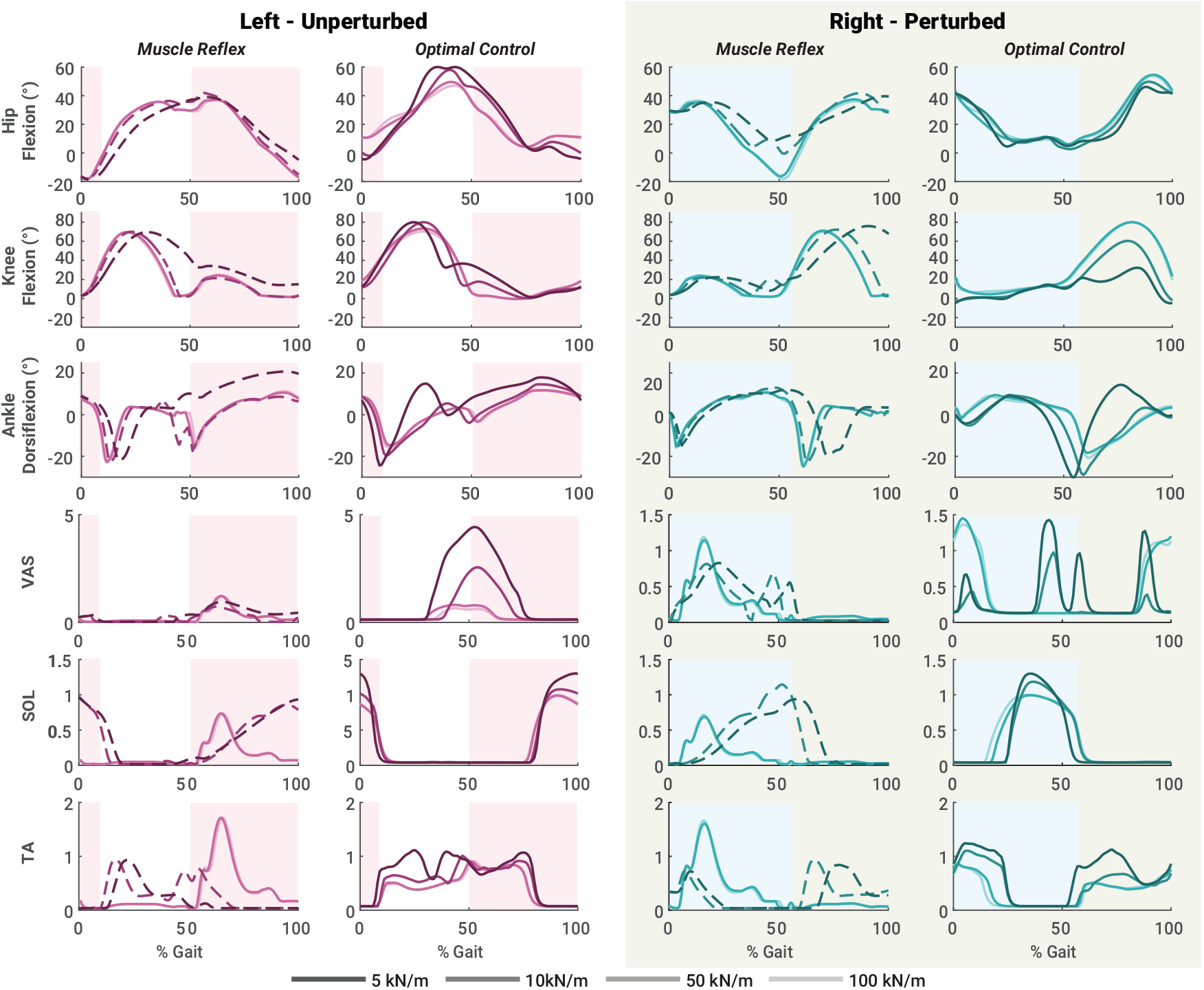
Bilateral hip, knee, and ankle kinematics and selected normalized muscle activations during one stride of steady-state walking with asymmetric foot-ground stiffness for both optimal control and muscle-reflex simulation methods. Darker lines correspond with lower stiffness on the perturbed side. Broken lines indicate muscle-reflex simulations that were not able to remain upright for the entire 30 step perturbation. Stride beginning and end is defined by heel-strike on the perturbed (right) side for all plots. Highlighted regions indicate stance phase for the corresponding side.

#### 2) Comparison of Muscle Activity

Fig. 3 illustrates mean muscle activations averaged across the gait cycle and normalized against the corresponding mean activations for the symmetrical, rigid ground condition. For the unperturbed side, the change of mean muscle activations with increasing stiffness value for two models matches up for all muscles except the GAS muscle. Optimal control simulations predicted a reduction in mean activation of GAS muscle for higher stiffness values, while the muscle-reflex model does not show a clear increasing or decreasing trend in GAS mean muscle activation. Optimal control simulations also predicted noticeably greater mean activation for the VAS muscle group than the muscle-reflex model. Mean muscle activations for the perturbed side have less agreement between the two models. Mean activation of GMAX, HAM and VAS muscles decreased with decreasing stiffness for optimal control simulations, while the muscle-reflex simulations show an opposite trend. Similarly, optimal control predicted a slight downward trend in mean SOL activation with decreasing stiffness, while muscle-reflex simulations estimated no change. Mean activation of the GAS muscle shows little change as ground stiffness decreases from the optimal control simulations, while the muscle-reflex model does not show linear change with higher values of perturbation stiffness. However, both models agree that mean TA muscle activity increases with decreasing ground stiffness.

**Fig. 3.**
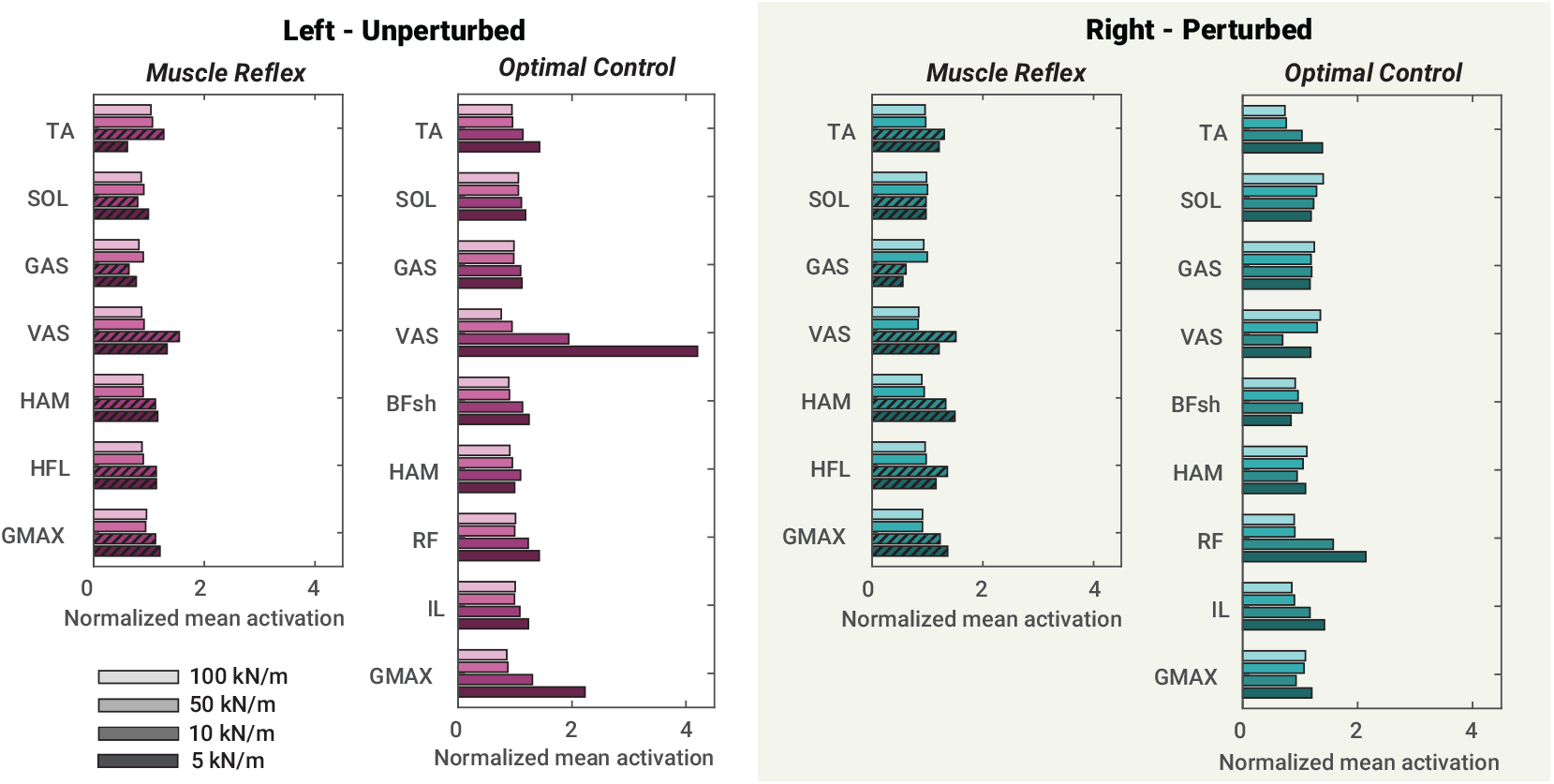
Bilateral normalized mean muscle activations averaged across one stride of steady-state walking with asymmetric foot-ground stiffness for both optimal control and muscle-reflex simulation methods. Darker bars correspond with lower stiffness on the perturbed side. Hatched bars indicate muscle-reflex simulations that were not able to remain upright for the entire 30 step perturbation.

Fig. 2 also shows the muscle activation profile throughout the gait cycle for steady-state walking for selected muscles: VAS, SOL, and TA. These muscles were chosen because experimental EMG data exists (SOL, TA) [7] or a major discrepancy between models was observed (VAS). For the unperturbed side of VAS muscle, optimal control simulations predict muscle activation peaks to be during the heel off phase, while muscle-reflex simulations estimate it to be slightly later during the gait cycle. As for the perturbed side of VAS muscle, the activation peak shifts from early stance to late stance as stiffness decreases with the optimal control method. However, with the muscle-reflex model, the muscle activation peak occurs during the mid-stance independent of the magnitude of the stiffness. As for the SOL muscle, the peak of muscle activation is earlier in the gait cycle for stiffness magnitudes of 50 kN/m and 100 kN/m and is later in the gait cycle for stiffness perturbation of 10 kN/m compared to optimal control simulations for both perturbed and unperturbed sides of the legs. The muscle-reflex model estimated significantly greater TA muscle activation peaks than optimal control simulations.

### B. Simulated vs. Experimental Results

#### 1) Comparison of Kinematics

Not only was the change of kinematic behavior for two models similar in response to a prolonged perturbation of asymmetric ground stiffness, it was similar to that observed experimentally in [7] in response to a single-step perturbation of asymmetric ground stiffness. The most notable difference between the simulation and experimental results was that neither model shows substantially increased dorsiflexion during swing on the unperturbed side for 10 kN/m stiffness, as observed in [7].

#### 2) Comparison of Muscle Activity

Likewise, the TA and SOL activation profiles resemble EMG data from a single-step perturbation in [7]. In simulations and experiments, TA and SOL activity increased on the unperturbed side with decreasing stiffness. The optimal control results more closely resemble the experimental EMG curves with respect to gait phase. The muscle reflex model produced spikes in TA activity during stance with high stiffness, which was not seen experimentally.

### C. Kinematic Aftereffects from Asymmetric Stiffness in the Muscle-Reflex Model

While the optimal control model did not afford analysis of how gait behavior changes over time, the muscle-reflex model did. As seen in Fig. 4, the gait kinematic patterns of the perturbed leg immediately and 30 strides after the removal of asymmetric stiffness perturbation were similar to those before the perturbation was applied. However, this was not the case for the unperturbed leg. While difference was subtle, peak knee flexion and peak ankle plantarflexion of the unperturbed leg were increased by ∼5°after the perturbation was applied compared to before the perturbation was applied.

**Fig. 4.**
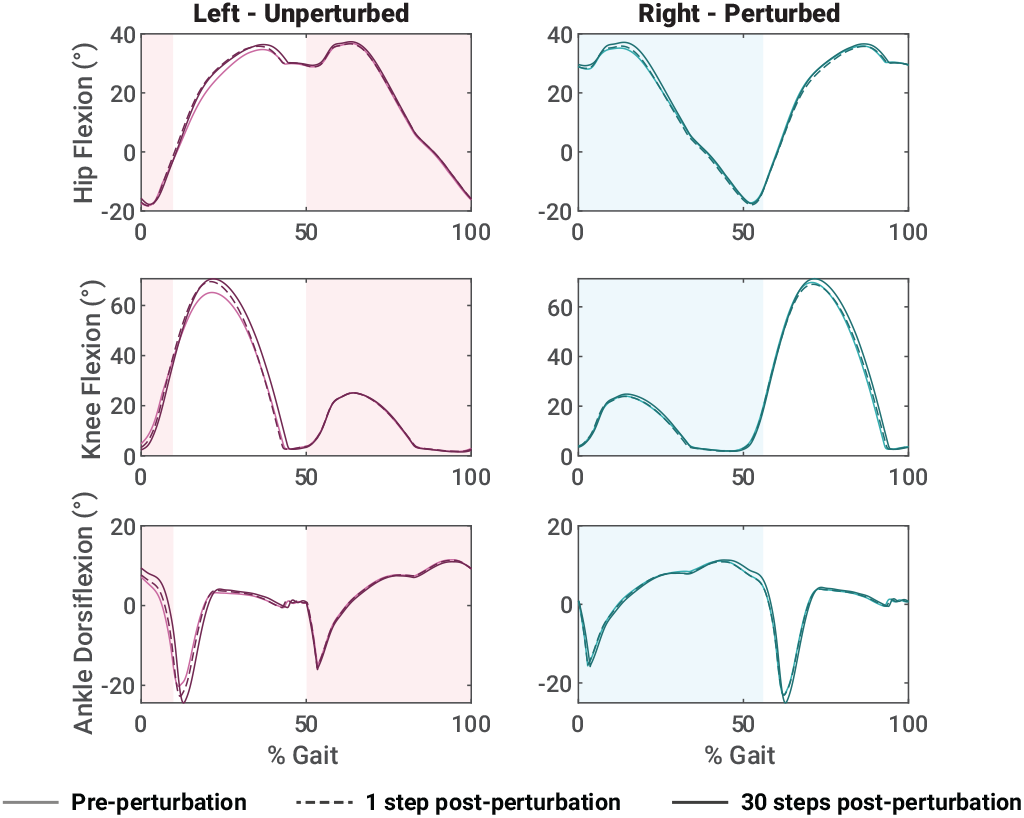
Muscle-reflex model joint kinematics pre- and post-perturbation. Light and dark lines indicate pre- and post-perturbation kinematics, respectively. Dashed and solid lines indicate immediate and 30-step after-effects, respectively.

### D. Effect of Asymmetric Stiffness and Damping on Muscle Effort in the Optimal Control Model

Fig. 5 illustrates the optimal control muscle effort cost function as a function of both asymmetric stiffness and damping. Damping had little influence on this metric for high stiffness, but had a large effect for stiffness values below 25 kN/m. An optimum stiffness for minimizing muscle effort can be observed around 10 kN/m for low damping, with the most extreme case at 5 Ns/m decreasing the cost function by 49% from the 1 MN/m baseline condition (note that this does not indicate a 49% reduction in metabolic cost). This low stiffness optimum disappears for damping values above 500 Ns/m, with muscle effort strictly increasing for lower stiffness. At 2000 Ns/m, the cost function is 55% greater for 10 kN/m than 1 MN/m, or 203% greater than the minimum effort condition at 10 kN/m stiffness and 5 Ns/m damping. Muscle effort increases relative to baseline for all damping values when stiffness falls to 7 kN/m or below.

**Fig. 5.**
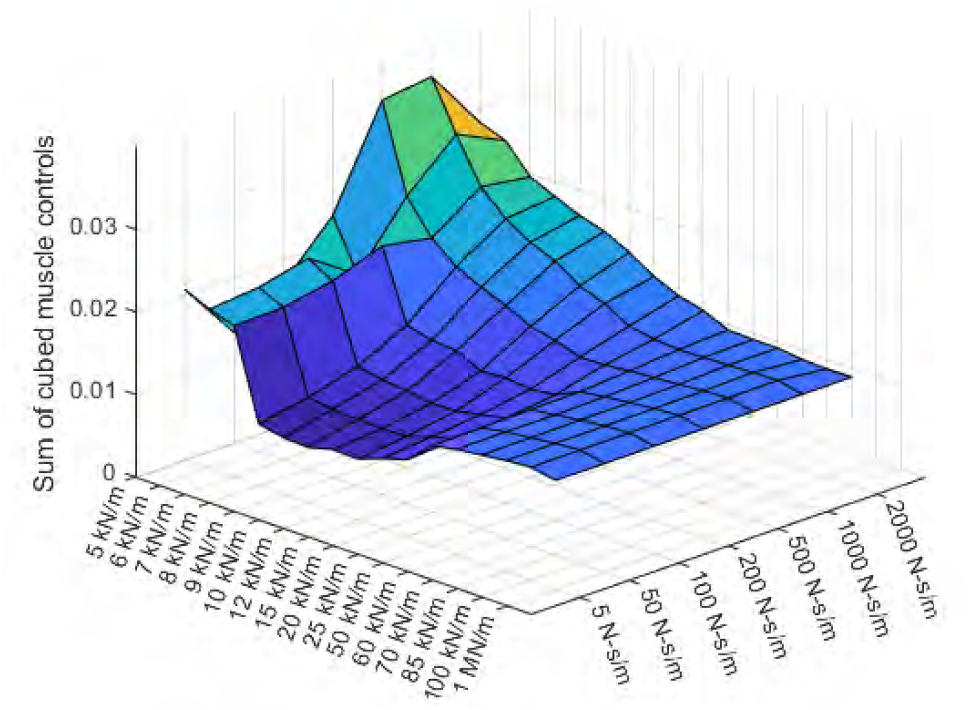
The optimal control muscle effort cost function value for the asymmetrical stiffness-damping landscape ranging from 5–1000 kN/m and 5–2000 Ns/m.

## IV. DISCUSSION

Overall, both models agree with the main trends observed in human experiments, in that contralateral TA activity and knee flexion increased as ground stiffness decreased for the opposite side leg. The optimal control model shows additional trends which agree with the experimental data: SOL activity, hip flexion, and ankle dorsiflexion also increased as opposite side ground stiffness decreased. This pattern is present despite the results corresponding with steady walking under the perturbed condition, rather than the single-step perturbation used experimentally. In general, however, these changes are not as apparent in simulation as they are in the experimental data except for lower ground stiffness values than were implemented experimentally. For the range of stiffnesses used in experiments, both models underestimate a trend which is specifically targeted as a desired rehabilitation outcome: the increase in contralateral ankle dorsiflexion and TA activity with decreasing stiffness.

The two approaches disagree with respect to the perturbed limb. The optimal control approach appears to utilize the reduced ground stiffness to reduce activity in some muscles on the perturbed side (SOL, VAS, HAM, GMAX). Conversely, muscle activity remains the same or strictly increases as stiffness decreases for the corresponding muscles in the muscle-reflex model. This discrepancy can partially be explained by the presence of additional muscles in the optimal control model. More broadly, it illustrates a fundamental difference in the two simulation methods. The optimal control method attempts to minimize muscle activity while meeting the task constraints, which tends to result in solutions that leverage the passive dynamics of gait, including the changed dynamics of introducing asymmetric ground stiffness. The muscle reflex model can be tuned to output an efficient gait for a fixed set of system dynamics, but is dependent on higher level control to exploit large changes in those dynamics, or even to maintain locomotion at all. The available experimental data for the perturbed side are limited to joint kinematics and few subjects [6], [25], and are not sufficient to validate these simulated patterns. However, these simulations suggest a useful framework to interpret results from future VST experiments.

The simulation methods have separate strengths and limitations. The muscle-reflex model struggles to stay upright for ground stiffness perturbations large enough to produce gait adjustments similar to experimentally observed adjustments. The model was even more sensitive to changes in damping and failed after minor alterations, preventing a systematic investigation. On the other hand, the optimal control model predicts that damping has a sizable effect on the relationship between muscle effort and asymmetrical ground stiffness, increasing overall muscle activity for lower stiffness as damping increased. More surprisingly, low damping simulations found an optimal stiffness range around 10 kN/m which decreased overall muscle effort relative to rigid ground, despite the stiffness being lowered asymmetrically.

Despite its robustness, the optimal control method as implemented with direct collocation does not simulate gait adaptations over time. Additionally, it does not represent a neuromotor control structure that can be used to model gait adaptations to changing foot-ground contact conditions, nor does it simulate interactions between supraspinal and peripheral neuromotor control. The muscle-reflex model is useful in this regard, because it can be used to simulate the behavior of the biomechanical system in absence of a central nervous system to provide evidence for the need for supraspinal control (or lack thereof) to recreate experimental behaviors. In this case, we observed signs of a post-perturbation aftereffect on the gait biomechanics despite the lack of a CNS, suggesting that experimentally observed aftereffects for VST experiments may not be fully explained by supraspinal adaptation, but may also reflect changes in the dynamical system response induced by the perturbation. The conditions which produced model failure were also the conditions which elicited the largest adjustments in the optimal control model, indicating that large changes in surface stiffness and damping may be necessary to elicit the neural activity required for rehabilitation.

While these two modeling approaches are useful, there still exists a need for a robust model which can simulate long-term adaptations to persistent foot-ground contact asymmetry. Methods such as single-shooting optimal control [26], [27] and reinforcement learning [28], [29] are better suited for simulating interactive perturbations and observing the change in behavior and may be appropriate for modeling the immediate aftermath of large stiffness or damping perturbations. There is also a need for new experiments using a VST to perform prolonged perturbations with additional EMG and kinematic data for both lower limbs, and to perform long term adaptation studies in general. Future simulation work will focus on investigating additional simulation methods.

